# BASOPHILS ACTIVATE PRURICEPTOR-LIKE VAGAL SENSORY NEURONS

**DOI:** 10.1101/2024.06.11.598517

**Authors:** Jo-Chiao Wang, Amin Reza Nikpoor, Théo Crosson, Eva Kaufmann, Moutih Rafei, Sébastien Talbot

## Abstract

Vagal sensory neurons convey sensations from internal organs along the vagus nerve to the brainstem. Pruriceptors are a subtype of neurons that transmit itch and induce pruritus. Despite extensive research on the molecular mechanisms of itch, studies focusing on pruriceptors in the vagal ganglia still need to be explored. In this study, we characterized vagal pruriceptor neurons by their responsiveness to pruritogens such as lysophosphatidic acid, *β*-alanine, chloroquine, and the cytokine oncostatin M. We discovered that lung-resident basophils produce oncostatin M and that its release can be induced by engagement of Fc*ε*RI*α*. Oncostatin M then sensitizes multiple populations of vagal sensory neurons, including Tac1^+^ and MrgprA3^+^ neurons in the jugular ganglia. Finally, we observed an increase in oncostatin M release in mice sensitized to the house dust mite *Dermatophagoides pteronyssinus* or to the fungal allergen *Alternaria alternata*, highlighting a novel mechanism through which basophils and vagal sensory neurons may communicate during type I hypersensitivity diseases such as allergic asthma.

## INTRODUCTION

Airway sensory innervation mostly originates from the jugular and nodose ganglia, projecting along the vagus nerve (1). These two mouse vagal sensory ganglia are anatomically fused and termed the jugular-nodose complex (JNC) (2). Like DRG neurons, jugular neurons are derived from the neural crest and express the lineage markers *Wnt1* and *Prdm12*, unlike nodose neurons, which originate from the neural placode and express the transcription factor *Phox2b* (*3, 4*). Pruriceptor neurons are primary afferent neurons that respond to itch-inducing agents known as pruritogens and generate itch signals transmitted to the somatosensory cortex through the spinal cord. While the mechanisms of itch generation in the skin have been extensively studied in recent years, research focusing on pruriceptors in vagal sensory neurons remains limited.

Basophils and mast cells differ in their maturation sites, and tissue localization but share several functional characteristics. These include activation upon the cross-linking of antigens with IgE, which is bound to the high-affinity IgE receptor, Fc*ε*RI*α* complex, on their plasma membranes. Upon activation, they release amine mediators such as histamine and serotonin and lipid mediators like cysteinyl leukotrienes and prostaglandins (5). Basophils are particularly noted for their high expression and rapid release of T_H_2 cytokines, including IL-4 and IL-13 (5). In severe asthmatic patients, basophils can react to bacterial products like Staphylococcus aureus enterotoxins (6) or exhibit elevated levels of receptors for alarmins, IL-25, IL-33, and TSLP (7).

Furthermore, basophil counts are associated with cold air-induced wheezing and cough (8) nighttime cough, and a history of asthma (9), suggesting a connection between basophils and neurogenic symptoms. Mechanistically, basophils promote acute itch by synthesizing and secreting LTC4 and IL-4 (10, 11). These mediators, in turn, act on pruriceptor neurons, thereby forming a skin basophil-neuron axis. It remains uncertain whether such basophil-neuron axis also exists in the airways and their functional role in the context of lung allergic diseases.

Recent advances in single-cell RNA sequencing have classified skin pruriceptor neurons into three main types, NP1, NP2, and NP3, based on their gene expression profiles. NP1 is characterized by the expression of Mas-related G protein-coupled receptor D (**MrgprD**) and lysophosphatidic acid receptor 3 (**LPAR3**). NP2 is defined by the expression of MrgprA3 and the GDNF family receptor alpha subunit 1 (**GFR*α*1**). NP3 is identified by the expression of cytokine receptors interleukin-31 receptor alpha (**IL-31R*α***) and oncostatin M receptor *β* (**OSMR**), as well as somatostatin (**SST**) and natriuretic peptide precursor B (BNP, encoded by ***Nppb***) (12). Recent single-cell RNA sequencing data indicate that mouse lung basophils highly express *Osm* and *Lif*, genes encoding the IL-6 family cytokines oncostatin M (OSM), leukemia inhibitory factor (LIF), and their signature cytokine IL-4 (13–15). Given that OSM has been implicated in mediating heat hypersensitivity and chronic itch (16, 17), and its cognate receptor, OSMR, is expressed on MrgprA3+ NP2 and *Nppb*+ NP3 pruriceptor neurons, we hypothesized that OSM could mediate basophils and vagal pruriceptor-like neurons crosstalk during allergic airway inflammation.

## RESULTS

Single-cell RNA sequencing data from Kupari et al. (refer to GSE124312 (18)) reveals that classic NP1 pruriceptor markers, *Lpar3* and *Mrgprd*, are expressed in one of the jugular clusters, JG2. In contrast, NP2 and NP3 markers such as *Il31ra*, *Mrgpra3*, *Osmr*, *Sst*, and *Nppb* are found in another jugular cluster, JG3 (**Fig 1A**). Notably, NP2-like and NP3-like jugular neurons are grouped into a single cluster, likely due to their gene expression similarities and low cell numbers, despite the mutually exclusive expression of *Mrgpra3* and *Il31ra* (**Fig 1A**). To further investigate whether these markers are co-expressed on the same neurons, we conducted an in-silico analysis of Zhao’s dataset (refer to GSE192987 (19)). We found that NP1-like vagal sensory neurons, which co-express *Lpar3* and *Mrgprd*, appear in one TRPV1-negative jugular (*Prdm12*^+^) cluster (**Fig 1B, SF 1A**).

**Figure 1:**
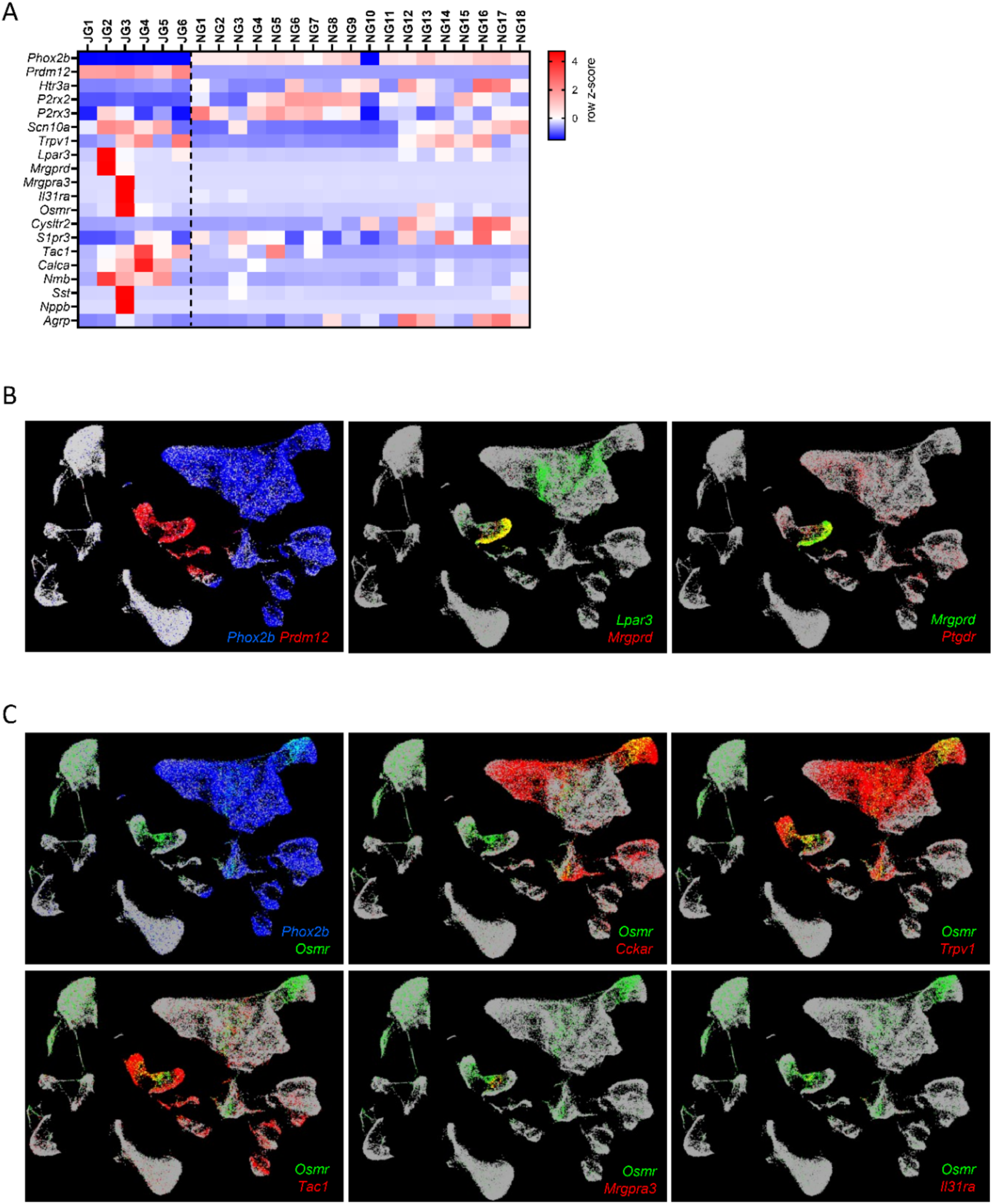
scRNAseq dataset revealing pruriceptor-like vagal sensory neurons. (**A**) *in-silico* analysis of the dataset GSE124312 generated a heatmap that shows the transcript expression levels of lineage-defining transcription factors (*Phox2b, Prdm12*), ion channels *(Htr3a, P2rx2, P2rx3, Scn10a, Trpv1*), pruriceptor markers (*Lpar3, Mrgprd, Mrgpra3, Il31ra, Osmr, Cysltr2, S1pr3*), and neuropeptides (*Tac1, Calca, Nmb, Sst, Nppb, Agrp*). (**B-C**) Uniform Manifold Approximation and Projection (UMAP) plots generated from the database GSE192987 feature pseudocolor highlighting of cells expressing specific genes, illustrating Phox2b-expressing placodal neurons and Prdm12-expressing neural crest neurons. Neurons labeled as NP1, which express Lpar3 and Mrgprd, are shown in plots (**B**). Meanwhile, cells expressing *Osmr*, which co-express *Trpv1, Tac1, Mrgpra3,* and *Il31ra*, are depicted in the plot (**C**). (**A**) The data are presented as a heatmap displaying each cluster’s z-scores of average gene expressions. Experimental details and cell clustering are defined in Kupari et al. (**B-C**). The data are shown as colored dots, and the cells are labeled with non-zero values in counts per ten thousand. The UMAP is derived from the database GSE192987, with experimental details outlined in Zhao et al.

*Osmr*-expressing neurons are present in both the jugular and nodose compartments. Nodose neurons express pruriceptor markers, including *Lpar3* and *Osmr*, but in clusters that do not co-express *Mrgprd*, *Mrgpra3*, or *Il31ra* (**Fig 1C**). Instead, nodose *Osmr*^+^ neurons express *Cckar1*, suggesting they are stomach-innervating neurons with intra-ganglionic laminar endings. Although rare, jugular Osmr^+^ neurons co-express classic NP2 and NP3 markers such as *Mrgpra3, Sst, Nppb,* and *Il31ra* (**Fig 1C**). It is also noted that jugular Osmr^+^ neurons can be further differentiated by the mutually exclusive expression of *Mrgpra3* and *Tac1* (**Fig 1C, SF 1B**). These findings suggest that putative pruriceptor-like vagal sensory neurons are primarily located in the jugular compartment of the mouse jugular-nodose complex.

Following identifying jugular pruriceptor-like vagal sensory neurons through transcriptomic analysis, we employ real-time calcium imaging to validate the functional expression of LPAR3 and MRGPRD using their respective ligands, xy-17 and *β*-alanine. Consistent with previous findings, neurons responding to xy-17 (LPAR3^+^) constitute half of the *β*-alanine-responding (MRGPRD^+^) neurons, with ∼5% of neurons responding to both xy-17 and *β*-alanine, and *∼*7% responding only to *β*-alanine (**Fig 2A**). These show minimal overlap with TRPV1^+^ (capsaicin-responding) neurons in the dorsal root ganglia (**Fig 2A**). In cultured jugular-nodose complex neurons, a smaller proportion of *β*-alanine-responsive neurons also respond to xy-17, with *∼*1% of neurons responding to both ligands and ∼3% responding to *β*-alanine (**Fig 2B**). Like DRG neurons, most *β*-alanine-responsive neurons in the JNC do not respond to capsaicin (**Fig 2B**).

**Figure 2:**
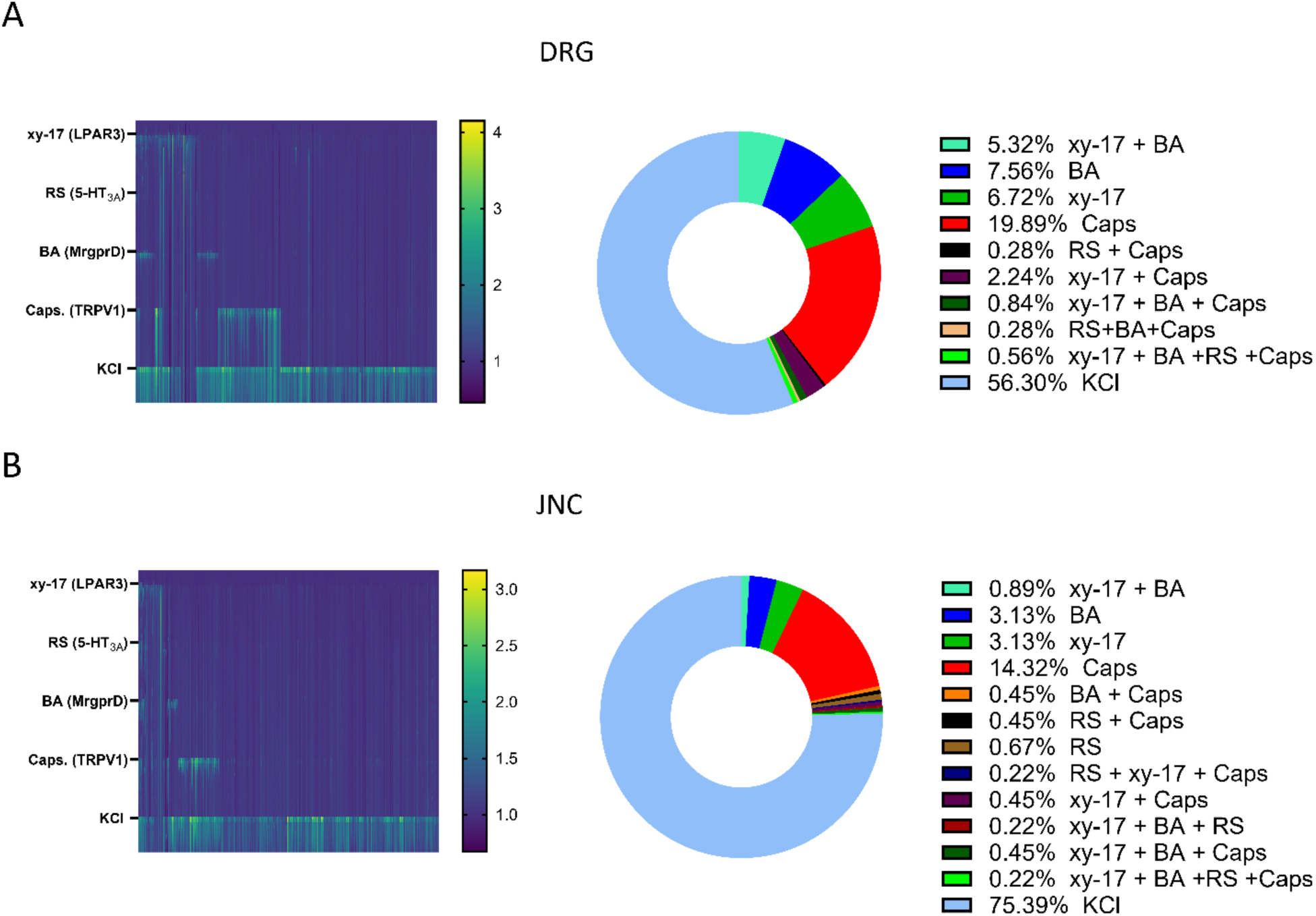
JNC neurons show markers of NP1 pruriceptor. (**A-B**) Calcium influx in cultured neurons from the dorsal root ganglia (**A**) and jugular-nodose complex (**B**) was measured in response to various agonists. Cells were sequentially treated with the LPAR3 agonist xy-17 (10 µM, from 60-70 sec), the 5-HT3A agonist RS56812 (RS; 1 µM from 360-380 sec), the MrgprD agonist *β*-alanine (BA; 1 mM from 660-680 sec), the TRPV1 agonist capsaicin (Caps; 1 µM from 960-970 sec), and KCl (100 mM from 1260-1280 sec). Calcium influx was visualized using the fluorescent dye fura-2 AM and is displayed as raster plots in the left panels. The proportions of neurons responding to different combinations of agonists are also shown.

Immgen data reveal lung basophils highly express IL-6 family cytokines (*Il6, Lif, Osm*) and their signature cytokine *Il4* (**SF 2A**). The human dataset from CELLxGENE indicates that human OSM expression can be detected in neutrophils and basophils (**SF 2B**). Since the expression of OSM by mouse basophils has not yet been reported, we will examine its expression in FACS-sorted cells from mouse lungs (**Fig 3A**), including eosinophils (Lin^-^ SiglecF^+^ c-kit^-^ Fc*ε*RI*α*^-^), alveolar macrophages (Lin^+^ SiglecF^+^), and T cells (Lin^+^ CD90.2^+^ CD49b^-^), and compare their *Osm* expression with that of basophils (Lin^-^ Fc*ε*RI*α*^+^ CD49b^+^). Despite being the least abundant cell population sorted (**Fig 3B**), basophils expressed the highest level of *Osm* (**Fig 3C**).

**Figure 3:**
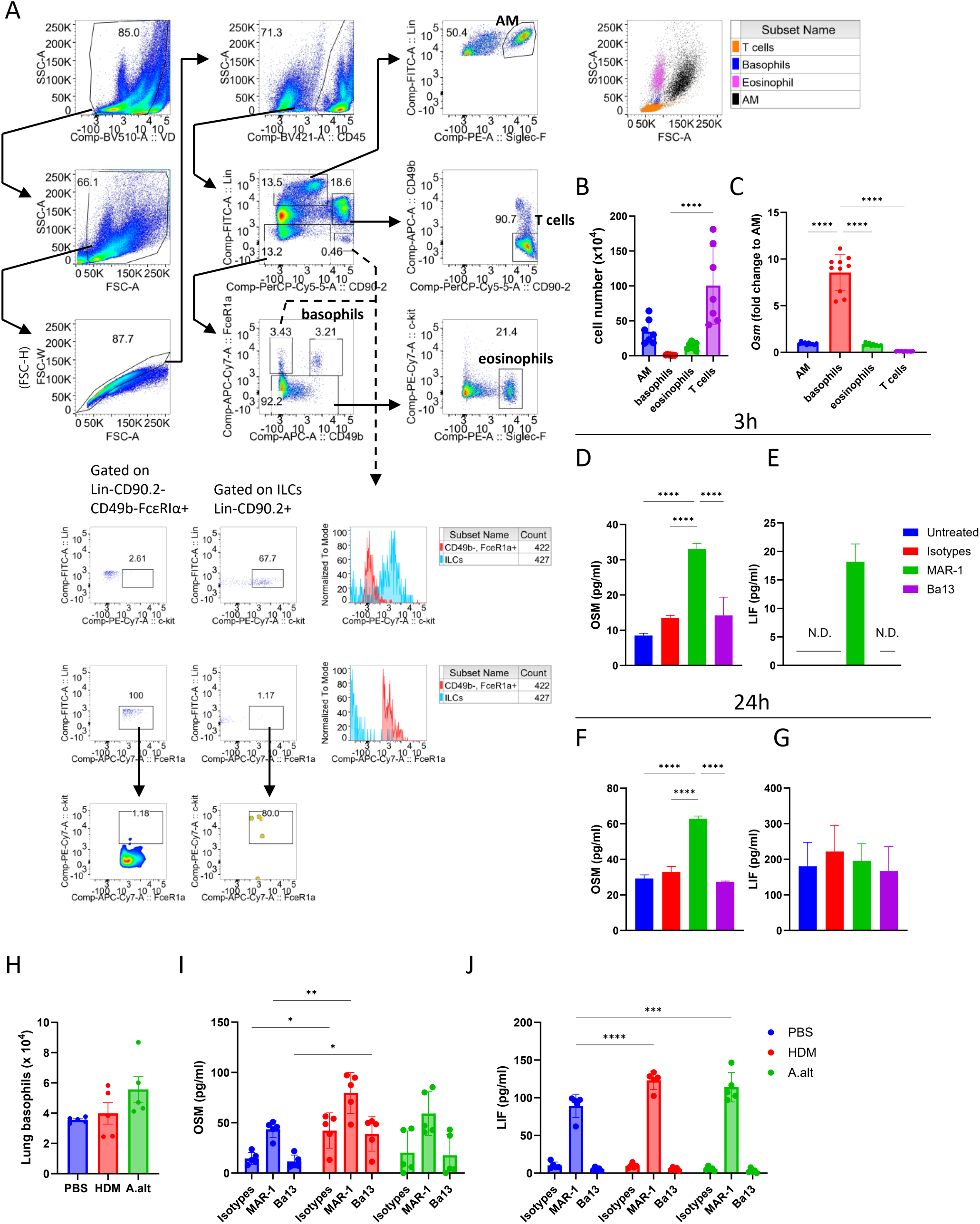
Lung basophils-release OSM upon Fc*ε*RI*α* engagement. (**A-C**) Gating strategy eosinophils (Lin^-^ SiglecF^+^ c-kit^-^ Fc*ε*RI*α*^-^), alveolar macrophages (AM, Lin^+^ SiglecF^+^), and T cells (Lin^+^ CD90.2^+^ CD49b^-^), basophils (Lin^-^ Fc*ε*RI*α*^+^ CD49b^+^), and c-kit-expressing cells under Lin^+^ CD90.2^+^ and Lin^+^ CD90.2^-^ CD49b^-^ Fc*ε*RI*α*^+^ gating. (**B-C**) Counts (**B**) and expression of *Osm* (**C**) in naïve mouse lung eosinophils (Lin^-^ SiglecF^+^ c-kit^-^ Fc*ε*RI*α*^-^), alveolar macrophages (AM, Lin^+^ SiglecF^+^), and T cells (Lin^+^ CD90.2^+^ CD49b^-^), basophils (Lin^-^ Fc*ε*RI*α*^+^ CD49b^+^) purified by FACS. (**D-G**) 10^6^ cells/well lung cells were cultured from C57BL6 mice and stimulated with MAR-1 (1 µg/mL), Ba13 (1 µg/mL), or a mix of isotype control antibodies. Levels of OSM (**D**, **F**) and LIF (**E, G**) in the supernatant of cultured lung cells were assessed by ELISA 3 hours (**D, E**) and 24 hours (**F, G**) post-stimulation. (**H-J)** 6-10 weeks old C57BL6 male and female mice were treated intranasally with PBS, house dust mite (HDM, 20 µg/dose), or Alternaria alternata (A.alt, 100 µg/dose) from day 0 to day four and challenged on day 7 to day 9. Basophil numbers were assessed by flow cytometry (**H**), while OSM (**I**) and LIF (**J**) levels in the supernatant of cultured lung cells (10^6^ cells/well) were assessed by ELISA at 3 hours post-stimulation with MAR-1 (1 µg/mL), Ba13 (1 µg/mL), or a mix of isotype control antibodies. (**A**) Representative FACS plot from one mouse. (**B**) Data are pooled from two independent experiments involving seven animals in each group. (**C**) Data are pooled from three independent experiments, with ten animals in each group. (**D-G**) Representative data from two independent experiments are shown. Lung cells were harvested and pooled from three animals, with three technical repeats for each group. (**H-J**) Representative data from two independent experiments, including five animals in each group and one technical repeat for cells from each animal. (**B-J**) Data are presented as FACS plots (**A**) or means ± SD (**B-J**). Statistical significance is indicated by *p*≤*0.05, **p*≤*0.01, ***p*≤*0.001, ****p*≤*0.0001.

Next, we sought to test whether activated basophils can release OSM. To test this, we stimulated cultured lung cells with antibody clones known to activate basophils, MAR-1 against Fc*ε*RI*α* and Ba13 against CD200R3. We measured OSM levels in the supernatant at 3- and 24-hours post-stimulation. MAR-1, but not unstimulated or isotype controls, significantly induced the release of OSM from cultured lung cells at both time points (**Fig 3D-E**). Additionally, MAR-1 drove the release of IL-4, the archetypical cytokine produced by basophils (**SF 3A**). MAR-1 also induced the release of LIF (another IL-6 family cytokine) at 3 hours post-stimulation (**Fig 3F**), but this was not sustained at 24 hours, as LIF levels accumulated over time in all conditions (**Fig 3G**). In contrast, Ba13 failed to induce the release of these cytokines (**Fig 3D-G**). Of note, eosinophils appear Fc*ε*RI*α*^-^ while ∼1% of Lin^-^ CD90.2^+^ Fc*ε*RI*α*^+^ were c-kit^+^ (**Fig 3A**). This data suggests that MAR-1-induced OSM release is mostly due to its action on lung basophils, as eosinophils do not express its receptor (Fc*ε*RI*α*^-^), and virtually no mast cells were present in our lung cell preparation.

We next assessed whether basal or induced OSM levels are altered in various mouse models of allergic airway inflammation (AAI), including those exposed to house dust mite *Dermatophagoides pteronyssinus* extract (HDM), *Alternaria alternata* medium (A.alt), or ovalbumin (OVA) in the presence or absence of fine particulate matter (FPM). Regardless of the types of infiltrated granulocytes (**SF 3B-E)**, we consistently observed increased expression of *Osm* in the lungs of mice with AAI (**SF 3G-F**). While there was no observed increase in total basophil numbers in homogenized lung tissue (**Fig 3H**), OSM release was found to be higher from cells of HDM-treated mice, both at basal levels (isotype control) and 3 hours post-MAR-1 treatment (**Fig 3I**). We also measured the levels of another IL-6 family cytokine, LIF, and found that it remained low at baseline across all groups. However, cells from mice treated with HDM and A.alt released higher levels of LIF upon MAR-1 stimulation (**Fig 3J**).

Given that the itch-amplifying cytokine oncostatin M counteracts the desensitization of capsaicin and histamine-induced inward currents, we sought to examine cultured vagal sensory neurons’ response to OSM. Initially, we discovered that OSM prevented TRPV1 desensitization at the population level in cultured DRG neurons (**SF 4A-B**), in contrast to JNC neurons (**SF 4C-D**). Next, we repeated this experiment for cultured JNC neurons using placode lineage reporter mice, Phox2b^cre^::Salsa6f^fl/wt^, in which cre-expressing cells are distinguished by their expression of the tdTomato-GCaMP6f fusion protein. First, we observed that neurons exposed to OSM — both Phox2b^+^ (identified as tdTomato-expressing) and Phox2b^-^ (non-tdTomato-expressing) — were protected from desensitization to the third capsaicin exposure (**Fig. 4A-D**). Next, to assess the capsaicin response of neural crest lineage neurons, we employ Tac1^cre^::Salsa6f^fl/wt^ mice since the Tac1^+^TRPV1^+^ neurons are restricted to the jugular compartment (**Fig. 1A, C**). Using these mice, we discovered that OSM-exposed Tac1^+^ neurons (identified as tdTomato-expressing) were protected from desensitization to the second capsaicin response (**Fig. 4E-F**), unlike Tac^-^ neurons (**Fig. 4G-H**). In all cases, the third response to capsaicin was preserved in OSM-exposed neurons (**Fig. 4A-H**).

**Figure 4:**
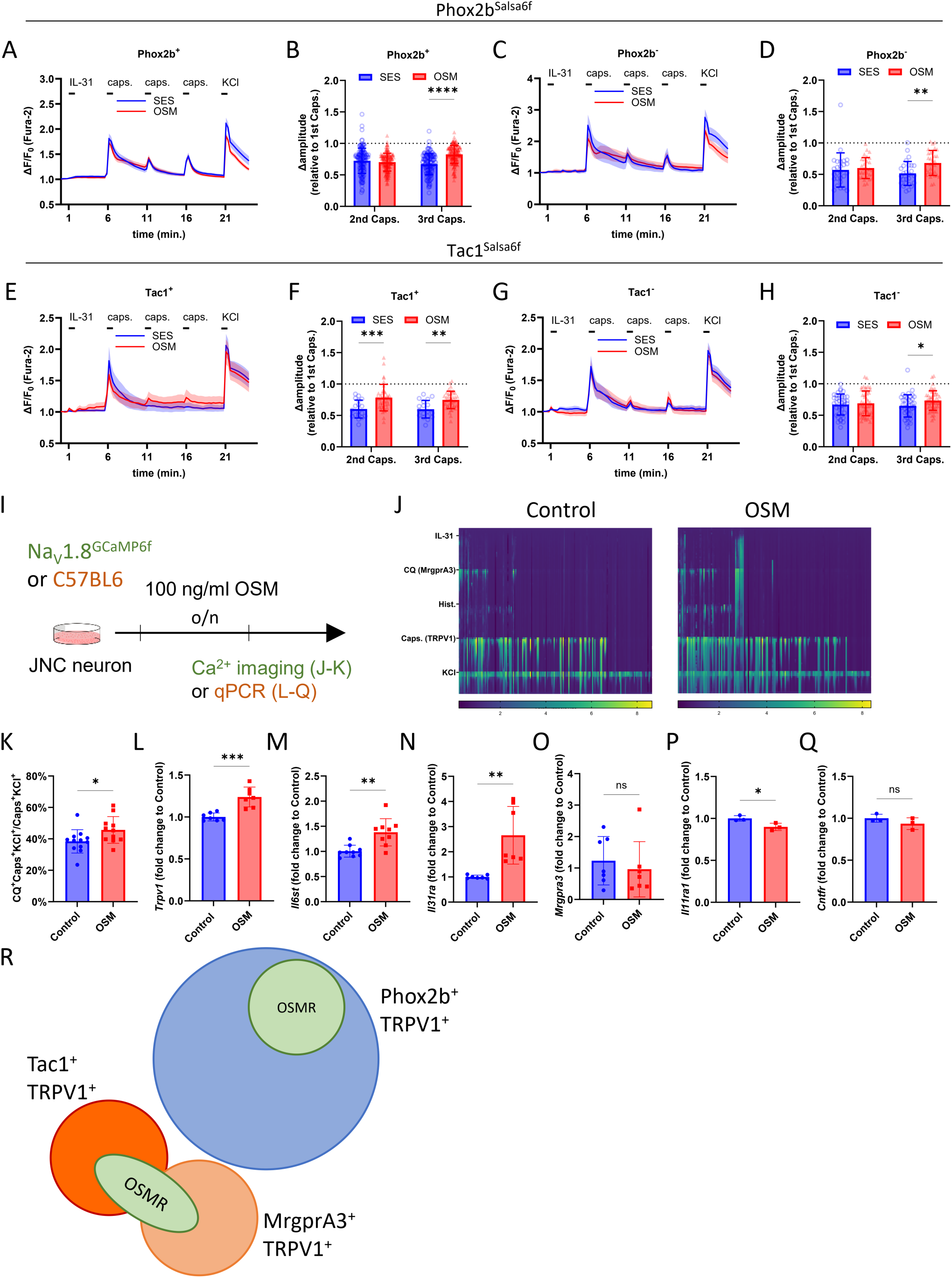
OSM prevents capsaicin-induced desensitization. (**A-H**) 6-12 weeks old male and female naïve Phox2b^cre^::Salsa6f^fl/wt^ mice (**A-D**) or Tac1^cre^::Salsa6f^fl/wt^ mice (**E-H**) were euthanized. JNC neurons were harvested, pooled, and cultured for 24 hours. After culturing, the cells were loaded with the calcium indicator Fura-2AM and sequentially stimulated with IL-31 (100 ng/mL; from 60 to 120 seconds), capsaicin (Caps; 100 nM; at 360-370 seconds, 660-670 seconds, and 960-970 seconds), and KCl (100 mM; from 1260 to 1280 seconds). Calcium flux was recorded throughout these stimulations. (**A, C, E, and G**) SES (blue) or 100 ng/ml OSM (red) was flown on the cells between the first and the second capsaicin exposure (420-660 sec). The second and third capsaicin responses of Phox2b^+^ (**B**; determined as tdTomato^+^), Phox2b^-^ (**D**; determined as tdTomato^-^), Tac1^+^ (**F**; determined as tdTomato^+^), and Tac1^-^ (**H**; determined as tdTomato^-^) neurons were normalized to the initial response (at 360-370 sec) of each neuron. (**I-K**) 6-12 weeks male and female naïve Na_V_1.8^cre^::GCaMP6f^fl/wt^ mice were euthanized, and JNC neurons were harvested, pooled, and cultured for 24h in the presence of vehicle or OSM (100 ng/mL). The cells were then sequentially stimulated with IL-31 (100 ng/ml; from 60 to 120 sec), MrgprA3 agonist chloroquine (CQ; 1mM 360-375 sec), histamine (Hist; 50 µM 660-720 sec), TRPV1 agonist capsaicin (Caps; 1 µM 960-970 sec), and KCl (40 mM 1260-1285 sec) and calcium flux recorded. (**K**) Proportions of CQ-responsive neurons among all capsaicin-responsive neurons. (**L-Q**) 6-12 weeks old male and female naïve C67BL6 mice were euthanized, JNC neurons harvested and cultured for 24h in the presence or absence of OSM (100 ng/mL). The cells were harvested, and gene expression of *Trpv1* (**L**), *Il6st* (**M**), *Il31ra (**N**), Mrgpra3* (**O**), *Il11ra1* (**P**), and *Cntfr* (**Q**) were evaluated by qPCR. (**A-R**) JNC neurons were harvested from 10 mice per experiment, pooled into ten dishes, and randomly assigned to either SES-treated or OSM-treated groups. One field of view was captured for each dish. (**A-B**) The analysis included n=138 capsaicin-responsive Phox2b^+^ SES-treated neurons and n=163 OSM-treated neurons. (**C-D**) Analysis of n=26 capsaicin-responsive Phox2b^-^ SES-treated neurons and n=30 OSM-treated neurons. (**E-F**) Analysis of n=18 capsaicin-responsive Tac1^+^ SES-treated neurons and n=38 OSM-treated neurons. (**G-H**) Analysis of n=40 capsaicin-responsive Tac^-^ SES-treated neurons and n=48 OSM-treated neurons. (**J-K**) JNC harvested from 10 mice per experiment was pooled into ten dishes and randomly divided into untreated or OSM-treated groups. One field of view was taken for each dish. (**J**) Representative data from 3 independent experiments: n=171 untreated neurons and n=152 OSM-treated neurons. (**K**) Pooled data from 3 independent experiments: n=12 fields of view in the control group and n=11 in the OSM-treated group. (**L-Q**) For cell culture, 104 cells/well were derived from 10-12 mice per experiment, with n=3-9 technical repeats from one experiment or pooled from 2-3 independent experiments. (**R**) An illustration showing the distribution of OSMR^+^ neurons in the mouse JNC. (**A**, **C**, **E**, **G**) Data are presented as means ± 95% CI of the maximum Fura-2AM (F/F_0_) fluorescence recorded every 15 seconds. (**B**, **D**, **F**, **H**, and **K-Q**) Data are presented as means ± SD. Statistical significance is indicated by *p*≤*0.05, **p*≤*0.01, ***p*≤*0.001, ****p*≤*0.0001.

Since jugular Osmr-expressing neurons can be further grouped into Mrgpr3A^+^ and Tac1^+^ subsets, we aimed to assess whether OSM affects responses to MrgprA3 stimulation with its agonist, chloroquine. Cultured Na_V_1.8^cre^::GCaMP6^fl/wt^ JNC neurons exposed to OSM overnight exhibited increased TRPV1^+^ neurons responding to chloroquine (**Fig 4J-K**). At the population level, cultured wild-type mouse JNC neurons exposed to OSM overnight displayed increased mRNA of *Trpv1* (**Fig 4L**), *Il6st* (**Fig 4M**), *Il31ra* (**Fig 4N**), but not *Mrgpra3*, *Il11ra1*, or *Cntfr* (**Fig 4O-Q**). mRNA of NP3 markers, *Htr1f* (**SF 4F**), *Hrh1* (**SF 4G**), *Cysltr2* (**SF 4H**), *S1pr1* (**SF 4I**), *Sst* (**SF 4J**), *Nppb* (**SF 4K**) and *Fstl1* (**SF 4L**), were not increased upon exposure of OSM, nor was *Tac1* (**SF 4M**). Our findings indicate that while OSM sensitizes placode-derived nodose neurons, its effect is more pronounced in neural crest-derived jugular and DRG neurons (**Fig 4R**).

Collectively, we present the first functional characterization of pruriceptor-like vagal sensory neurons present in the mouse jugular-nodose complex (JNC). This characterization includes NP1-like neurons that respond to xy-17 and *β*-alanine and NP2/3-like neurons that can be sensitized by oncostatin M (OSM)—a cytokine produced by lung basophils, with levels that increase during allergic airway inflammation.

## DISCUSSION

The existence and the overall role of dedicated pruriceptor neurons in the autonomic nerve systems still need to be tested. Here, we utilized a combination of lineage reporter lines to investigate subtypes of vagal sensory neurons originating from the placode or neural crest, focusing on their ability to detect pruritogenic mediators. Specifically, we characterized the responsiveness of vagal pruriceptor neurons to pruritogens such as lysophosphatidic acid, *β*-alanine, chloroquine, and the cytokine oncostatin M. We discovered that oncostatin M is produced by lung-resident basophils, with its release triggered by Fc*ε*RI*α* engagement. Furthermore, oncostatin M prevents the desensitization of nociceptors across various populations of vagal sensory neurons, notably in Tac1^+^TRPV1^+^ and MrgprA3^+^ neurons within the jugular ganglia. Additionally, we observed an increase in oncostatin M release in house dust mite (HDM)-sensitized mice. These data reveal a potential novel crosstalk between basophils and vagal neurons, which could exacerbate type I hypersensitivity diseases such as allergic asthma.

### Jugular and nodose ganglia

In mice, the jugular and nodose ganglia are anatomically fused. While agonists of P2X_2/3_ (20) and protease-activated receptor 1 (PAR-1) (21) can selectively activate nodose neurons, a specific activator for jugular neurons needs to be better defined. The expression of conventional nociceptor markers such as TRPV1, TRPA1, or TAC1 is not exclusive to jugular vagal sensory neurons. However, scRNAseq data indicates that the expressions of MrgprA3 and MrgprD are unique to jugular vagal sensory neurons, offering a potential new method to study the activity of jugular neurons.

### Vagal pruriceptors

Accumulated studies using guinea pigs suggested that activating jugular C-fiber neurons, which innervate the large extrapulmonary airways, evokes a cough reflex (22). Prostaglandin D2 (PGD2) is elevated in asthmatics and triggers strong bronchoconstriction (23–25). Since jugular MrgprD^+^ neurons express the PGD2 receptor, their bronchoconstriction effect is likely mediated through these neurons. This effect is similar to that of BAM8-22 and NPFF, activating pruriceptor features in airway-innervating vagal sensory neurons (26). Finally, combining the cysteinyl leukotriene receptor antagonist montelukast with fluticasone helps control cough-variant asthma (27).

### Lysophosphatidic acid

Lysophosphatidic acid (LPA) can be synthesized through extracellular autotaxin (28). Intracellular LPA induces itch by activating receptors such as LPAR5, TRPV1, or TRPA1 (29). LPA is present in bronchoalveolar lavage fluid collected from allergic or asthmatic patients following a provocative airway allergen challenge (30). It enhances T_H_2 inflammation and barrier integrity by interacting with eosinophils and epithelial cells (31). When introduced intravenously, LPA activates the carotid body, activating the vagal nerve and subsequent bronchoconstriction (32). This pathway was exaggerated in an ovalbumin (OVA)-induced rat asthmatic model (33). Given xy-17-induced calcium flux in cultured vagal sensory neurons, LPAR3-expressing NP1-like vagal sensory neurons will likely respond to LPA in concert with the carotid body during asthma attacks. This highlights the potential multifaceted role of LPA in exacerbating asthma and mediating sensory responses through vagal sensory neurons.

### Oncostatin M

Oncostatin M (OSM), a member of the IL-6 cytokine family, typically signals through heterodimer receptors consisting of OSMR and gp130. OSM can also bind to the LIF receptor (LIFR) in rats and humans (34, 35). Elevated levels of OSM are observed in idiopathic pulmonary fibrosis (IPF) (36), chronic obstructive pulmonary disease (COPD) (37), asthma with incomplete reversible airflow (38), coronavirus disease (COVID19) (39, 40). The cellular source of OSM has been a topic of debate. While some studies point to neutrophils (41), monocytes, or T cells (16), single-cell RNA sequencing datasets suggest that basophils highly express Osm. Here, we corroborate these transcriptomic data, showing that basophils express *Osm* mRNA and release it upon the engagement of Fc*ε*RI*α*. The absence of high-affinity IgE receptors in mouse eosinophils and the scarcity of mast cells in lung parenchyma mean that Fc*ε*RI*α*-based stimulation is predominantly directed to basophils. While in humans, *Osm* is highly detected in neutrophils in the CELLxGENE single-cell RNA sequencing datasets, this observation could be influenced by the scarcity of basophils in the sampled tissues. Additionally, the normalization methods used in this database may not adequately account for bias introduced by batch effects, necessitating further research to confirm OSM production from human basophils. Moreover, there are notable differences between mice and humans concerning the basophil-OSM-preceptor axis, including variations in receptor-binding affinities, which could impact the biological implications of this pathway in different species.

### Neuron-basophil crosstalk

A basophil activation test can be a convenient surrogate for traditional skin allergy tests in allergic patients (42). Not only does this test overcome the challenge of measuring short-lived unbound IgE in the serum(43), but the correlation between basophil activation state and disease severity highlights their role in the pathophysiology of allergy. In mouse models, allergen-induced acute itch typically involves the mast cell-histamine pathway. However, in the context of atopic dermatitis-associated inflammation, this pathway becomes redundant, and a previously unrecognized basophil-leukotriene axis takes precedence in mediating acute itch flares (10). This finding points to a novel basophil-neuronal circuit that could play a role in various neuroimmune processes. In agreement with this, we found elevated levels of OSM and LIF in mice with allergic airway inflammation, suggesting a reinforced basophil-neuronal crosstalk at the inflammatory state.

Basophils have traditionally been viewed as a circulating population recruited to the lungs during T_H_2-mediated inflammation (44, 45). However, Cohen et al. demonstrated that basophils are present in the lungs from birth, where they interact with epithelial cells to facilitate the maturation of local macrophages into alveolar macrophages (13). Despite the relatively small number (∼20,000 cells) of lung resident basophils, our data indicate that these cells express high levels of IL-4 and IL-6 family cytokines and can readily secrete these cytokines upon Fc*ε*RI activation. Given that i) no significant increase in lung basophil numbers is observed in allergic mice, and ii) cytokines were released by basophils cultured from naïve mice, we infer that these cytokines are predominantly released from lung resident basophils. These findings underscore the emerging role of tissue-resident basophils in inflammatory processes. Furthermore, considering that a broader range of sensory neurons expresses the leukemia inhibitory factor receptor (LIFR) compared to those expressing OSMR, the ability of OSM to signal through LIFR may allow it to influence a wider array of neuronal types beyond just preceptor neurons. We also observed that LIF production could be induced by Fc*ε*RI*α* engagement in mice, suggesting the potential for mouse basophils to interact with a broader spectrum of neurons beyond those expressing OSMR.

### OSM maintains pruriceptor-like vagal sensory neuron activity

Capsaicin, the active component of chili peppers, is a well-known agonist of the transient receptor potential vanilloid 1 (TRPV1) channel, predominantly expressed in sensory neurons (46). Activation of TRPV1 by capsaicin leads to an influx of calcium ions, causing the sensory neurons to fire and convey signals perceived as heat or pain (46). Repeated or prolonged exposure to capsaicin can induce a phenomenon known as desensitization, where the sensitivity of TRPV1 to subsequent stimulation is reduced (47). This desensitization occurs through calcium-dependent dephosphorylation, receptor internalization, and decreased channel responsiveness (48). Studies suggest that desensitization is a protective mechanism to prevent overactivation of sensory neurons, which can be beneficial in managing pain (48). This mechanism has therapeutic implications, as capsaicin patches and creams are used in clinical settings to exploit TRPV1 desensitization for pain relief in conditions such as neuropathic pain and osteoarthritis.

On the other hand, compounds that prevent TRPV1 desensitization are likely to drive nociceptor neurons, the cells expressing TRPV1, activity and, therefore, amplify allergic inflammation (49–52). Notably, protein kinase complexes A and C (PKA and PKC) play a crucial role in maintaining responses to capsaicin. While nodose OSMR^+^ neurons co-express *Prkaca*, which encodes the PKA catalytic subunit *α* (**SF 1C**), jugular OSMR^+^ neurons co-express *Prkca*, encoding PKC-*α* (**SF 1D**). Notably, *Prkcq*, encoding PKC-*θ*, is specifically expressed in jugular MrgprD-expressing neurons (**SF 1E-F**), indicating further specialization within these sensory pathways. These findings suggest distinct pathways in preventing capsaicin desensitization by OSM in different lineages of OSMR^+^ vagal sensory neurons.

## CONCLUSION

Our findings illustrate that preceptor-like vagal sensory neurons are predominantly jugular neurons that can interact with basophil-produced OSM. This interplay might be important for exacerbating neurogenic symptoms in allergic asthma.

## MATERIALS AND METHODS

### Animals and experimental procedures

Animals and experimental procedures. All procedures involving animals were conducted in accordance with the guidelines of the Canadian Council on Animal Care (CCAC) and the Queen’s University Animal Care Committee (UACC). Mice were housed in individually ventilated cages with access to water and subjected to 12-hour light cycles; food was available ad libitum. C57BL6/J (000664), Phox2b-Cre (016223), Tac1-IRES2-Cre-D (021877), NaV1.8-Cre (036564), Ai95(RCL-GCaMP6f)-D (C57BL/6J) (028865), and LSL-Salsa6f (031968) were obtained from the Jackson Laboratory and bred in-house. Lineage-specific reporter mice were produced by breeding the cre lines with Ai95(RCL-GCaMP6f)-D or LSL-Salsa6f lines, facilitating calcium imaging experiments.

### Ovalbumin-induced lung inflammation

For the ovalbumin (OVA)-induced allergic airway inflammation model, C57BL6 mice were sensitized by intraperitoneally injecting an emulsion of grade V OVA (200 µg/dose; Sigma-Aldrich A5503) and Imject® Alum (1 mg/dose; Thermo Fisher 77161) on days 0 and 7, followed by intranasal challenges with OVA (50 µg/dose) with or without FPM (20 µg/dose) from day 14 to 16. Control mice were sensitized but not challenged. The mice were sacrificed on day 17 to harvest tissues.

### House dust mite-induced lung inflammation

C57BL6 mice received daily intranasal injections of house dust mite extract (CiteQ 02.01.85, 20 µg/dose) from day 0 to 5 as sensitization, and from day 8 to 10 as challenges. PBS was injected into control mice. Mice were sacrificed on day 11 following anesthesia with intraperitoneally injected urethane (2 g/Kg body weight; Sigma Aldrich 94300) to harvest tissues.

### *Alternaria alternata*-induced lung inflammation

C57BL6 mice received daily intranasal injections of *Alternaria alternata* media (CiteQ 09.01.26, 100 µg/dose) from day 0 to 5 as sensitization, and from day 8 to 10 as challenges. PBS was injected into control mice. Mice were sacrificed on day 11 following anesthesia with intraperitoneally injected urethane (2 g/Kg body weight; Sigma Aldrich 94300) to harvest tissues.

### *In-silico* analysis of gene expression by vagal sensory neurons

Data were extracted from the supplementary materials of Kupari et al., with clusters defined according to the original publication (18). Raw data from GSE192987 were obtained from the National Center for Biotechnology Information (NCBI) website and re-plotted with the following quality control filtering parameters: number of genes per cell greater than 200, number of genes per cell less than 8000, and percentage of mitochondrial genes less than 5% per cell.

### Neuron culture

JNC and DRG were extracted and dissociated as previously described (53, 54). Briefly, JNC was collected following exsanguination, while DRG was collected after the decapitation of anesthetized mice. Ganglia were placed into a digestion buffer containing 1 mg/ml (325 U/ml) collagenase type 4 (Worthington LS004189), 2 mg/ml (1.8 U/ml) Dispase II (Sigma 04942078001), and 250 µg/ml (735.25 U/ml) DNase I (Sigma 11284932001), prepared in supplemented DMEM media, and incubated at 37°C for 60 minutes. Mechanical dissociation was performed by pipetting the digested tissue with pipette tips of decreasing diameter and finishing with 25-gauge needles, followed by density gradient centrifugation with 150 mg/ml bovine serum albumin (BSA; Hyclone SH30574.02; PBS solution) over a PBS layer. Cells were seeded onto glass-bottom dishes (ibidi 81218) pre-coated with 50 µg/ml laminin (Sigma L2020) and 100 µg/ml poly-D-lysine (Sigma P6407) and cultured overnight in supplemented Neurobasal-A media before recording calcium imaging.

### Bronchoalveolar lavage fluid (BALF) and lung tissue harvest

Bronchoalveolar lavage was performed on mice anesthetized as previously described, with incisions made to the trachea. The mice were lavage twice with 1 ml of PBS or FACS buffer (2% FBS and 1 mM EDTA in PBS) using a Surflo ETFE IV Catheter 20G x 1” (Terumo Medical Products SR-OX2025CA). The collected lavage fluid was centrifuged at 350 x G for 6.5 minutes. The supernatant was collected for ELISA, and the cell pellets were resuspended, subjected to RBC lysis (Cytek TNB-4300-L100 or Gibco A1049201), and stained for surface markers for flow cytometry analysis. Lungs were harvested following a diaphragm incision and transcardial perfusion with 10 ml of PBS, then minced with razor blades, and collected into TRIzol™ Reagent (Invitrogen 15596026) for RNA extraction or into a digestion buffer (1.6 mg/ml collagenase type 4 and 100 µg/ml DNase I in supplemented DMEM) to prepare a single-cell suspension. This suspension was obtained through 45 minutes of enzymatic digestion at 37°C, with mechanical dissociation using 18-gauge needles performed midway through incubation (30 minutes), followed by filtering through a 70 µm nylon mesh, and RBC lysis. For flow cytometry analysis or fluorescence-activated cell sorting (FACS), cells were resuspended in FACS buffer; for in vitro stimulation, cells were resuspended in FBS-supplemented DMEM, seeded into 96-well plates, and cultured at 37°C with 5% CO_2_ for the specified time before supernatant collection.

### ELISA

Cytokine levels in supernatant collected from BALF or cell culture were determined with commercially available ELISA kits from Biolegend, R&D systems, Cusabio, and MyBioSource.

### *In vitro* stimulation of lung cells

Dissociated lung cells were cultured in 96-well tissue culture plates treated to support cell growth, with a density of 1x10^6 cells per well. The cells were treated with 1 µg/ml MAR-1 (Biolegend), 1 µg/ml Ba13 (Biolegend), or a mixture of corresponding isotype antibodies. Supernatants were collected at 3 hours and 24 hours post-stimulation in the initial experiment. The 3-hour time point was chosen to compare cytokine release from cells treated under different conditions, as the basal release of cytokines is lower at this time point.

### Fluorescence-activated cell sorting (FACS)

Single-cell suspensions derived from BALF, or lung samples were stained with Ghost Dye Violet 510 (Cytek. 13-0870-T100) and antibody cocktails in PBS at 4°C for 30 minutes, followed by fixation with 10% neutral buffered formalin (Sigma Aldrich. HT501128) at room temperature for 15 minutes prior to flow cytometry data acquisition. For assessing eosinophil and neutrophil infiltration, BALF cells were stained with fluorochrome-conjugated antibodies targeting CD45 (30-F11), CD90.2 (53-2.1), CD11b (M1/70), CD11c (N418), Ly6C (HK1.4), Ly6G (1A8), and Siglec-F (1RNM44N). For basophil sorting, cells from BALF or lung were stained with antibodies against CD45 (30-F11), CD90.2 (53-2.1), CD49b (DX5), FcεRIα (MAR-1), c-kit (2B8), Siglec-F (1RNM44N), and lineage markers including CD3ε (145-2C11), CD19 (1D3/CD19), NK1.1 (PK136), and CD11c (N418). These antibodies were sourced from Biolegend or Thermo Fisher Scientific. For FACS, cells were resuspended in FACS buffer post-staining without fixation, filtered again using Falcon® Round-Bottom Tubes with a Cell Strainer Cap (35 µm strainer, Corning 38030), and analyzed. Data acquisition was performed using a FACS Canto II (BD Biosciences) or CytoFLEX S (Beckman Coulter) flow cytometer. Cell sorting was conducted on a BD FACSAria III or BD FACSAria Fusion.

### Calcium imaging

Cultured neurons from C57BL6, Phox2b^cre^::Salsa6f^fl/wt^, or Tac1^cre^::Salsa6f^fl/wt^ were loaded with the calcium indicator dye, 5 µM fura-2 AM (Cayman Chemical Company 34993), and incubated at 37°C for 40 minutes. After incubation, they were washed four times with standard external solution (SES; Boston BioProducts C-3030F) and subsequently imaged in SES. Neurons from NaV1.8-GCaMP6f mice were washed with SES and immediately used for imaging. In experiments using Salsa6f reporter lines, the fluorescent signal from GCaMP6f was shown to completely coincide with tdTomato+ cells (data not shown). The data recorded with signals from Fura-2 were used for further analysis. Agonists were diluted in SES and administered through a ValveLink8.2 system (AutomateScientific) with 250 µm Perfusion Pencil® tips (Automate Scientific), facilitated by Macro Recorder (Barbells Media, Germany). SES was continuously flowed during intervals between drug injections to wash out the drugs. Imaging for Fura-2 experiments was conducted using an S Plan Fluor ELWD 20X objective lens (NIKON) to enhance UV light passage, while GcaMP6f experiments utilized an S Plan Fluor LWD 20X lens (NIKON) for improved resolution. Images were captured at 3 or 4-second intervals using pco.edge 4.2 LT (Excelitas Technologies), Prime 95B (Teledyne Photometrics), or Orca Flash 4.0 v2 (Hamamatsu Photonics) sCMOS cameras. All imaging was performed on ECLIPSE Ti2 Inverted Microscopes (NIKON). Regions of interest (ROI) were manually delineated on NIS-Elements software (NIKON), and the F340/F380 ratio or GFP measurements were exported to Excel (Microsoft) for further analysis. Data were condensed into a maximum value every 15 seconds for all analyses.

### Real-time quantitative PCR (qRT-PCR)

Sorted cells, freshly minced lung tissues, or digested lung single-cell suspensions were lysed using TRIzol Reagent and stored at -80°C prior to RNA extraction. RNA from sorted cells was extracted using PureLink RNA Micro Scale Kits (ThermoFisher 12183016), while RNA from lung tissues or lung cell suspensions was extracted using E.Z.N.A.® Total RNA Kit I (Omega Bio-tek® R6834). All RNA extractions were conducted according to the manufacturer’s instructions, following phenol-chloroform phase-based purification and mixing with equal volumes of isopropanol. cDNA synthesis was carried out using SuperScript VILO Master Mix (Invitrogen 11755050) with 1-2 µg of RNA template for each reaction. Quantitative polymerase chain reaction (qPCR) was performed using PowerUp SYBR Green Master Mix (Applied Biosystems A25742), 50-100 ng of cDNA templates, and 200 nM of respective primers on a Mic qPCR Cycler (Bio Molecular Systems) or CFX Opus Real-Time PCR System (Bio-Rad Laboratories).

### Data availability

Information and raw data are available from the lead contact upon reasonable request.

### Statistics

P *values ≤ 0.05* were considered statistically significant. One-way ANOVA, two-way ANOVA, and Student t-tests were performed using GraphPad Prism. DESeq2 and Seurat analysis and statistics were performed using RStudio.

### Replicates

Replicates (n) are described in the figure legends and represent the number of animals for *in vivo* data. For *in vitro* data, replicates can either be culture wells or dishes, animals, fields-of-view (microscopy), or neurons (calcium microscopy), but always include different preparations from different animals to ensure biological reproducibility.

## ACKNOWLEDGEMENTS

ST work is supported by the Canadian Institutes of Health Research (CIHR; 407016, 461274, 461275), Canadian Foundation for Innovation (44135), Canadian Cancer Society Emerging Scholar Research Grant (708096), Knut and Alice Wallenberg Foundation (KAW 2021.0141, KAW 2022.0327), Swedish Research Council (2022-01661), Natural Sciences and Engineering Research Council of Canada (RGPIN-2019-06824), and NIH/NIDCR (R01DE032712). Salary support for JCW was provided by the Fonds de recherche du Québec – Santé (FRQS), the Canadian Allergy, Asthma, and Immunology Foundation, Asthma Canada (CAAIF), and CIHR (Institute of Circulatory and Respiratory Health). MR is supported by the Canadian Institutes of Health Research project grant (PJT-186233) and EK is supported by the Canada Research Chair program.

---

**Supplementary Figure 1:**
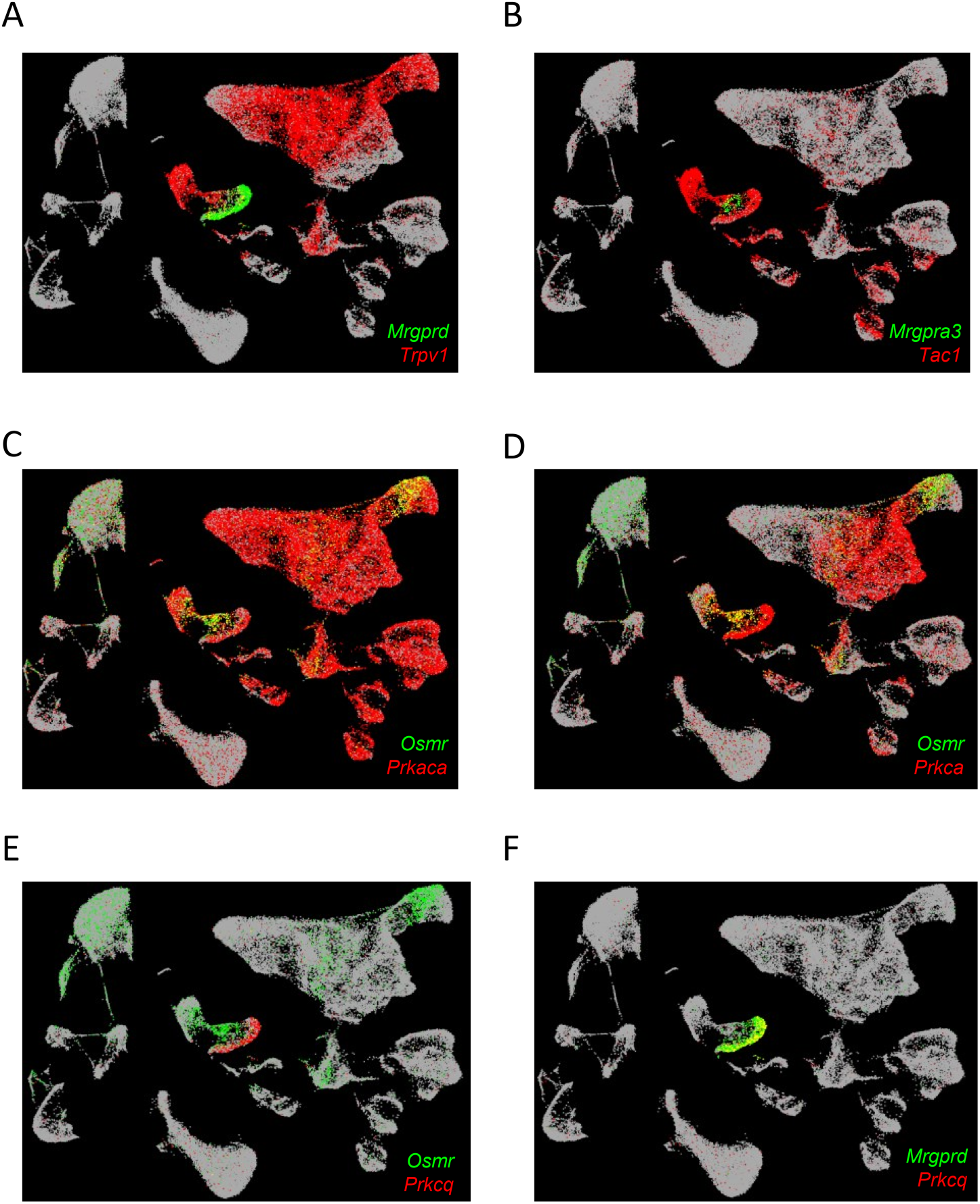
*Mrgprd*, *Prkca* and *Prkcq* in JNC neurons. *in silico* analysis of the GSE192987 single-cell RNA sequencing data reveals specific gene expression patterns in jugular neurons. *Mrgprd*^+^ jugular neurons do not express *Trpv1* (**A**), and *Mrgpra3*^+^ neurons do not express *Tac1* (**B**). Further analysis highlights the transcript expression patterns of *Prkaca* (**C**), *Prkca* (**D**), and *Prkcq* (**E-F**), along with those of *Osmr* (**C-E**) and *Mrgprd* (**F**). (**A-F**) Data are presented as pseudocolors, representing cells with non-zero counts per ten thousand. The UMAP plots were generated using data from the database GSE192987. Experimental details were outlined in Zhao et al.

**Supplementary Figure 2:**
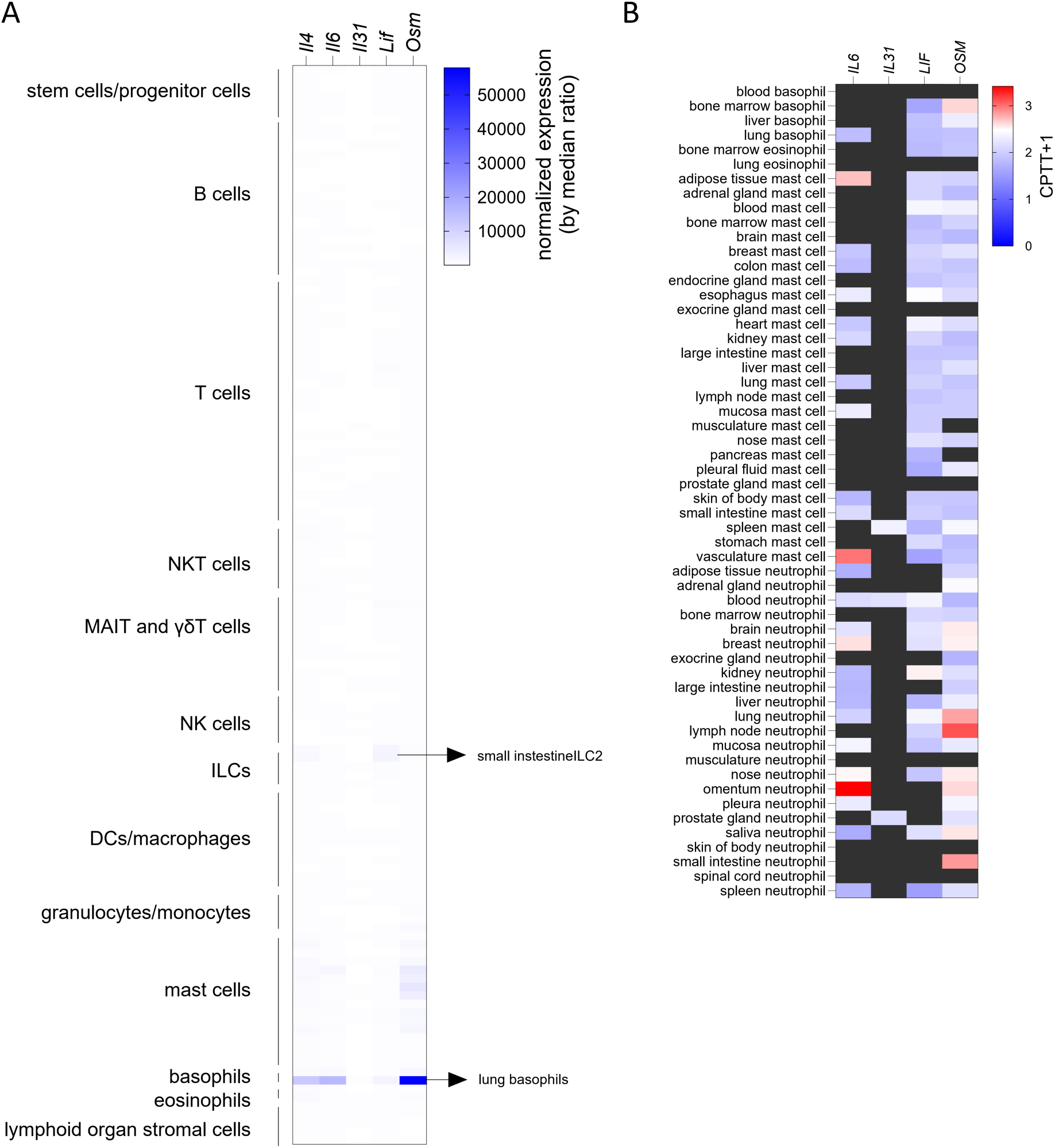
Mouse basophils and human neutrophils express OSM. *in silico* analysis of OSM expression in mouse (Immgen; **A**) and human (CELLxGENE; **B**) immune cells reveal its expression in mouse basophils (**A**) and human neutrophils (**B**). (**A**) Data are presented as normalized gene expression using the median of ratios method. Detailed in (1).(**B**), Data are presented as cluster average of gene expression normalized by counts per ten thousand + 1 of genes having a non-zero value and detailed in (2).

**Supplementary Figure 3:**
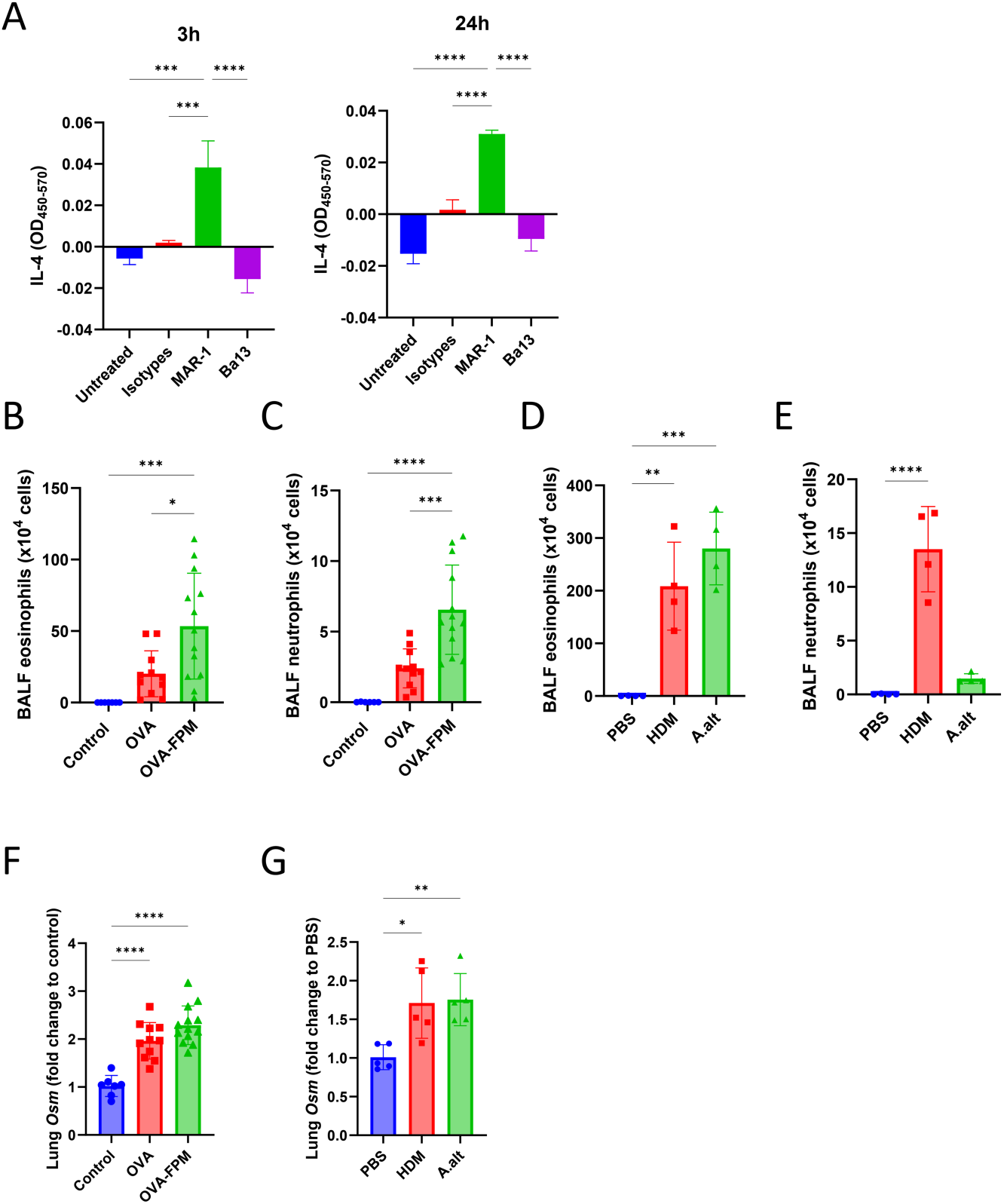
Lung basophils-release IL-4 upon Fc*ε*RI engagement. (**A**) 10^6^ cells/well lung cells were cultured from naïve C57BL6 and stimulated with MAR-1 (1 µg/mL), Ba13 (1 µg/mL), or a mix of isotype control antibodies. IL-4 levels measured in the supernatant of cultured lung cells were assessed by ELISA 3 hours and 24 hours post-stimulation with MAR-1 (1 µg/mL), Ba13 (1 µg/mL), or a mix of isotype control antibodies. (**B-G**) 6-10 weeks old male and female C57BL6 mice were sensitized with an intraperitoneal injection of an emulsion containing OVA (200 µg/dose) and aluminum hydroxide (1 mg/dose) on days 0 and 7. Subsequently, they were challenged intranasally with OVA (50 µg/dose) with or without fine particulate matter (FPM, 20 µg/dose) (**B, C, and F**). Other groups of 8-week-old male and female naïve C57BL6 mice were treated intranasally with house dust mite (HDM, 20 µg/dose) or Alternaria alternata (A.alt, 100 µg/dose) from day 0 to day four and challenged on day 7 to day 9 (**D, E, and G**). Upon sacrifice, bronchoalveolar lavage fluid (BALF) was harvested, and eosinophil (**B, D**) and neutrophil (**C, E**) infiltration were assessed by flow cytometry and *Osm* expression measured in whole lung lysates (**F, G**) by qPCR. (**A**) Data from one experiment. Lung cells were harvested and pooled from three animals, with n=3 technical repeats for each group. (**B, C,** and **F**) Pooled data from three independent experiments, with n=7 in the control group, n=11 in the OVA group, and n=13 in the OVA-FPM group. (**D, E**) Representative data from two independent experiments, with n=4 in each group. (**G**) Representative data from two independent experiments, with n=5 in each group. Data are presented as means ± SD, with statistical significance indicated as *p*≤*0.05, **p*≤*0.01, ***p*≤*0.001, ****p*≤*0.0001.

**Supplementary Figure 4:**
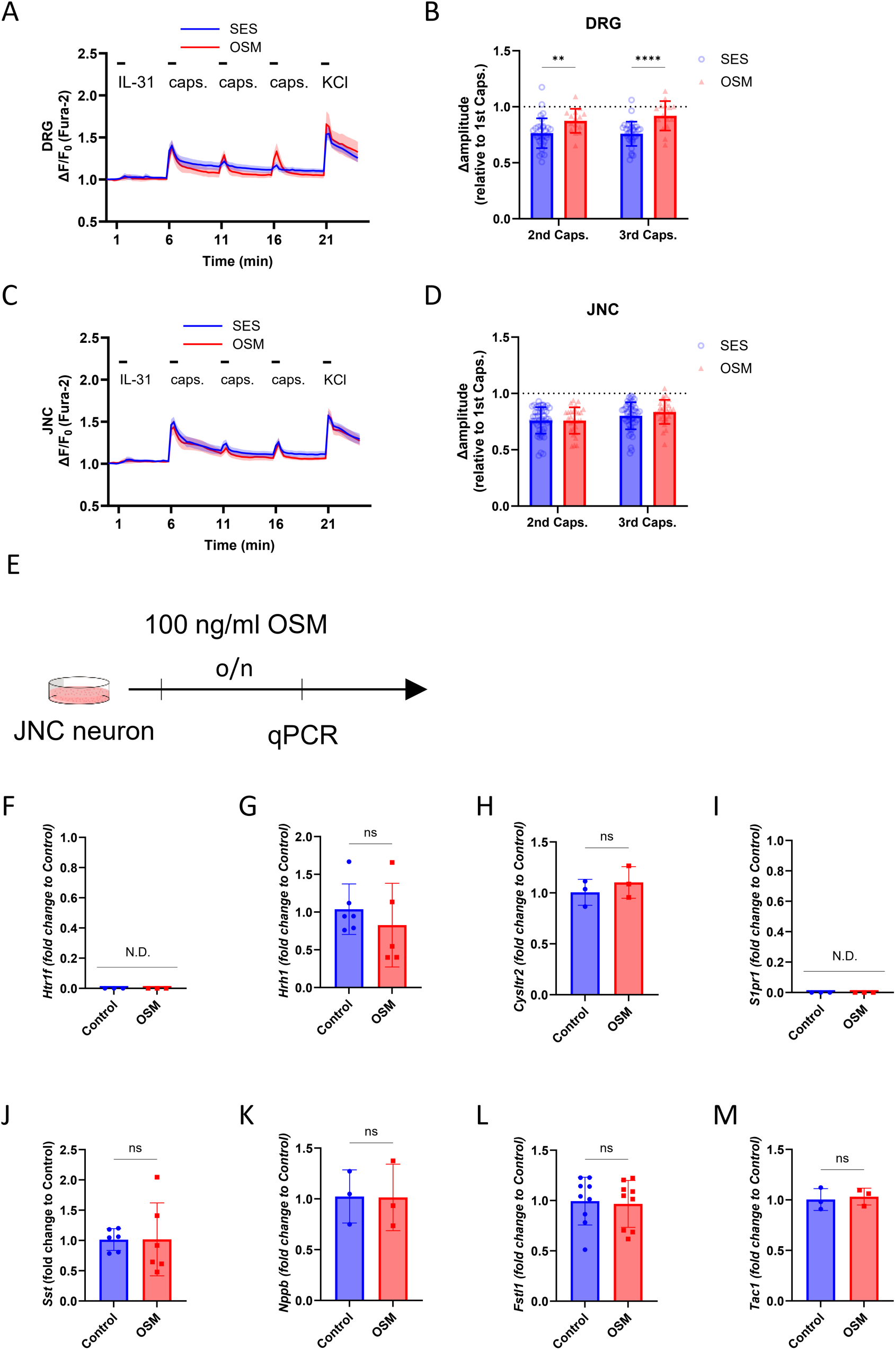
OSM prevents capsaicin-induced desensitization in DRG neurons. (**A-D**) 6-12 weeks old male and female naïve C57Bl6 mice were euthanized, and DRG (**A-B**) or JNC (**C-D**) neurons were harvested and cultured for 24h. The cells were then loaded with Fura-2AM and sequentially stimulated with IL-31 (100 ng/mL; from *60 to* 120 sec), capsaicin (Caps; 100 nM; from 360 to 370, 660 to 670, and 960 to 970 sec), and KCl (100 mM; from 1260 to 1280 *sec*) and calcium flux recorded. (**A and C**) SES (blue) or 100 ng/ml OSM (red) was flown on the cells between the first and the second capsaicin exposure (420-660 sec). (**B and D**) The second and third capsaicin responses were normalized to each individual neuron’s initial response (at 360-370 sec). (**E-M**) 6-12 weeks old male and female naïve C57BL6 mice were euthanized, JNC neurons harvested and cultured for 24h in the presence of a vehicle or OSM (100 ng/mL). The cells were harvested, and transcript expressions of *Htr1f* (**F**), *Hrh1* (**G**), *Cysltr2* (**H**), *S1pr1* (**I**), *Sst* (**J**), *Nppb* (**K**), *Fstl1* (**L**), and *Tac1* (**M**) were assessed. (**A-D**) JNCs harvested from 10 mice per experiment were pooled into 10 dishes, and DRG from each animal was cultured in 4 dishes. Dishes were randomly assigned to either the SES or OSM-treated group. One field of view was taken for each dish. Representative data from 2 independent experiments are shown. n=33 capsaicin-responsive SES-treated and n=15 OSM-treated DRG neurons were analyzed (**A**), and n=49 capsaicin-responsive SES-treated and n=28 OSM-treated JNC neurons were analyzed (**C-D**). (**A** and **C**) Data are presented as means ± 95% CI of the maximum Fura-2AM (F/F_0_) fluorescence every 15 seconds. (**B, D**, and **F-M**) Data are presented as means ± SD. Statistical significance is indicated as *p*≤*0.05, **p*≤*0.01, ***p*≤*0.001, ****p*≤*0.0001.

## SUPPLEMENTARY MATERIALS

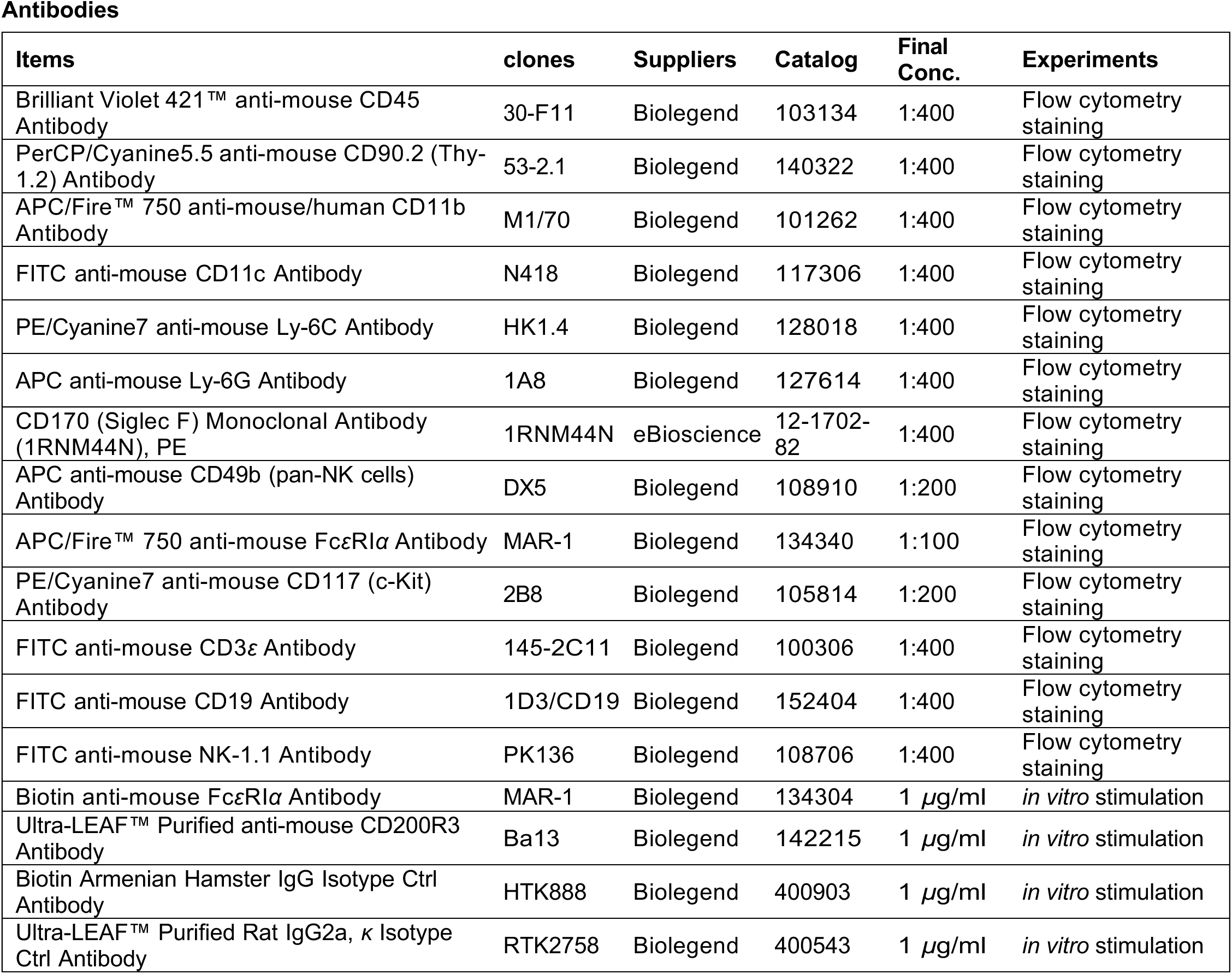
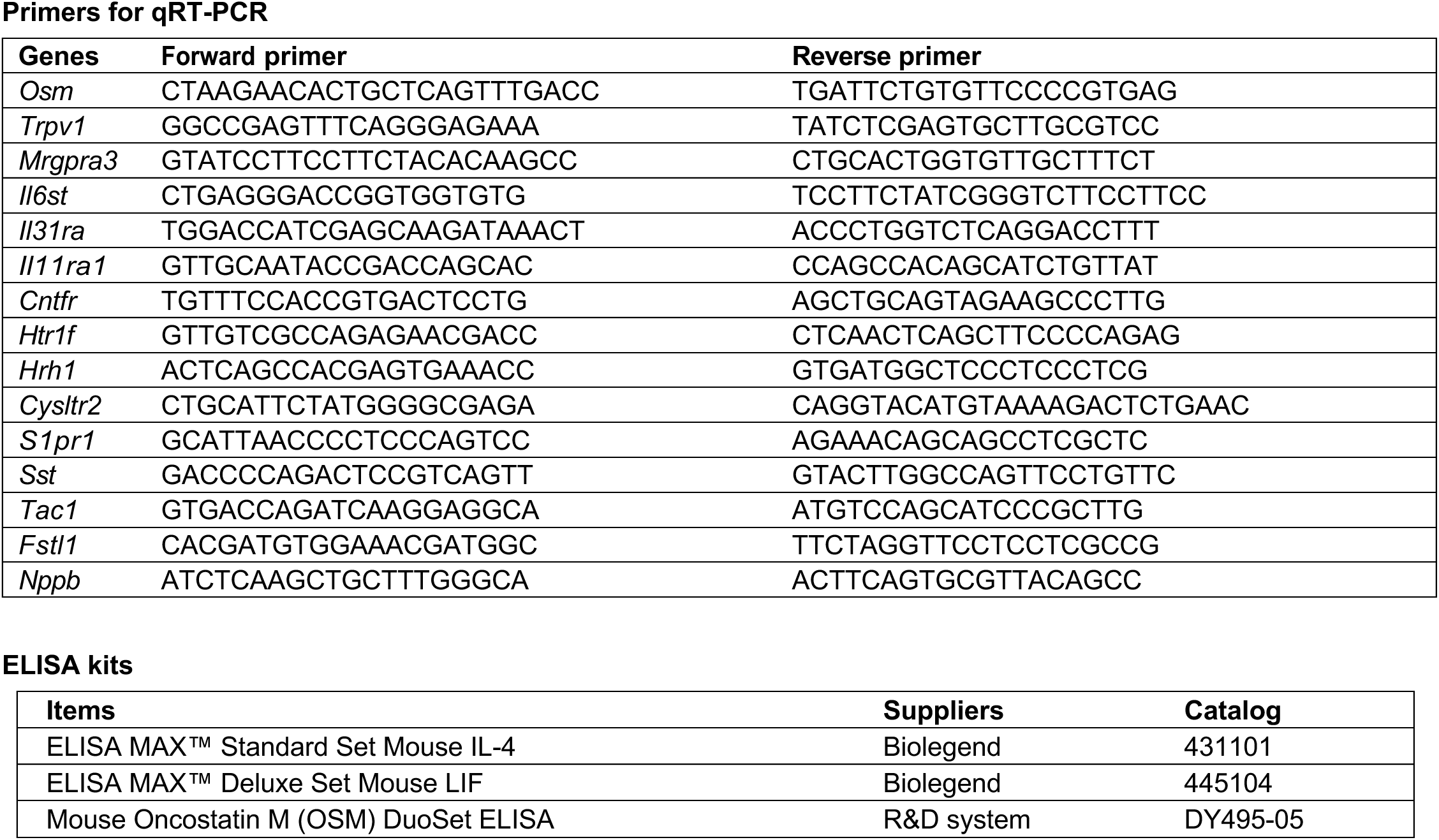

### Buffers, Cell culture media, and supplements

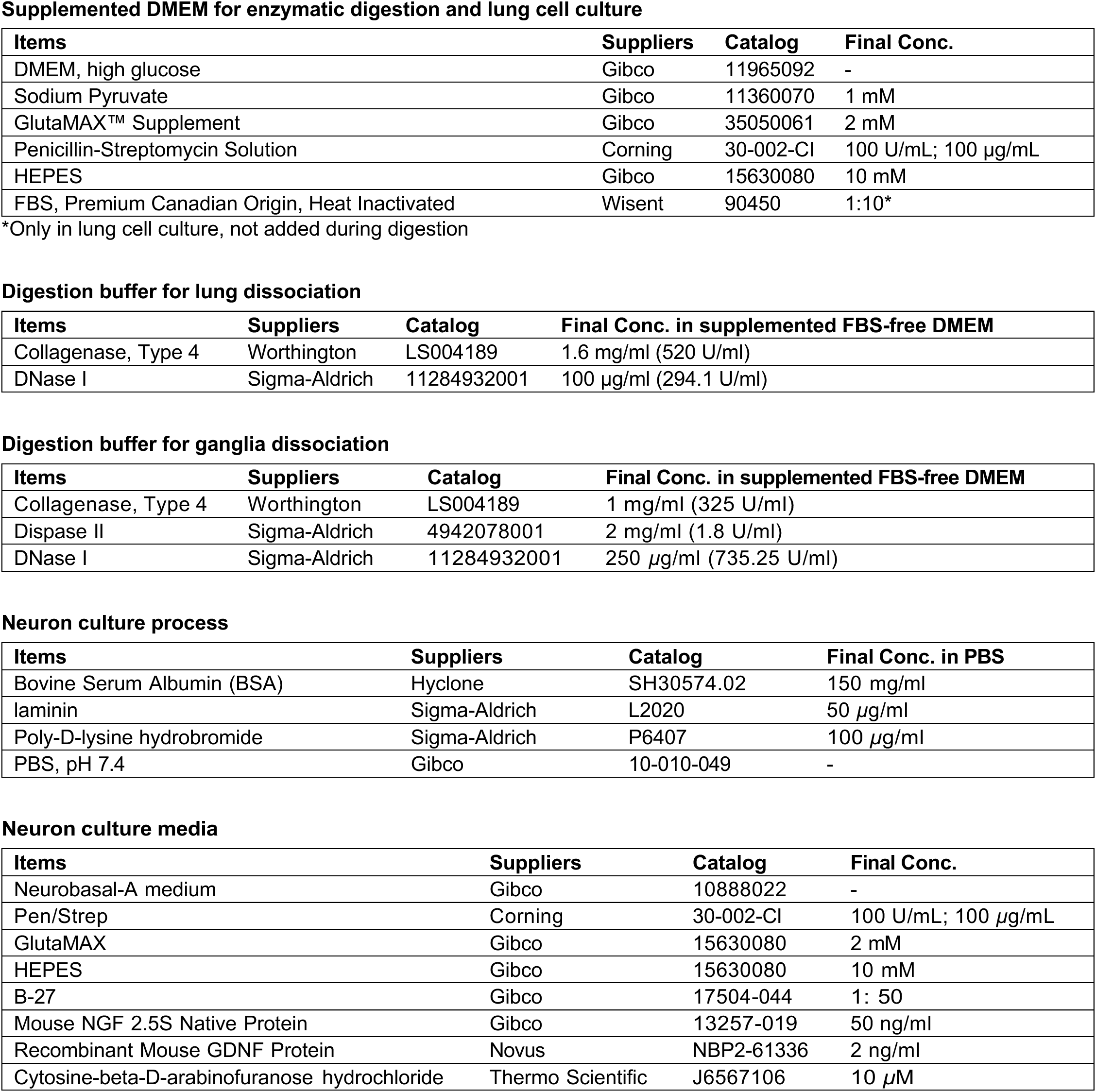
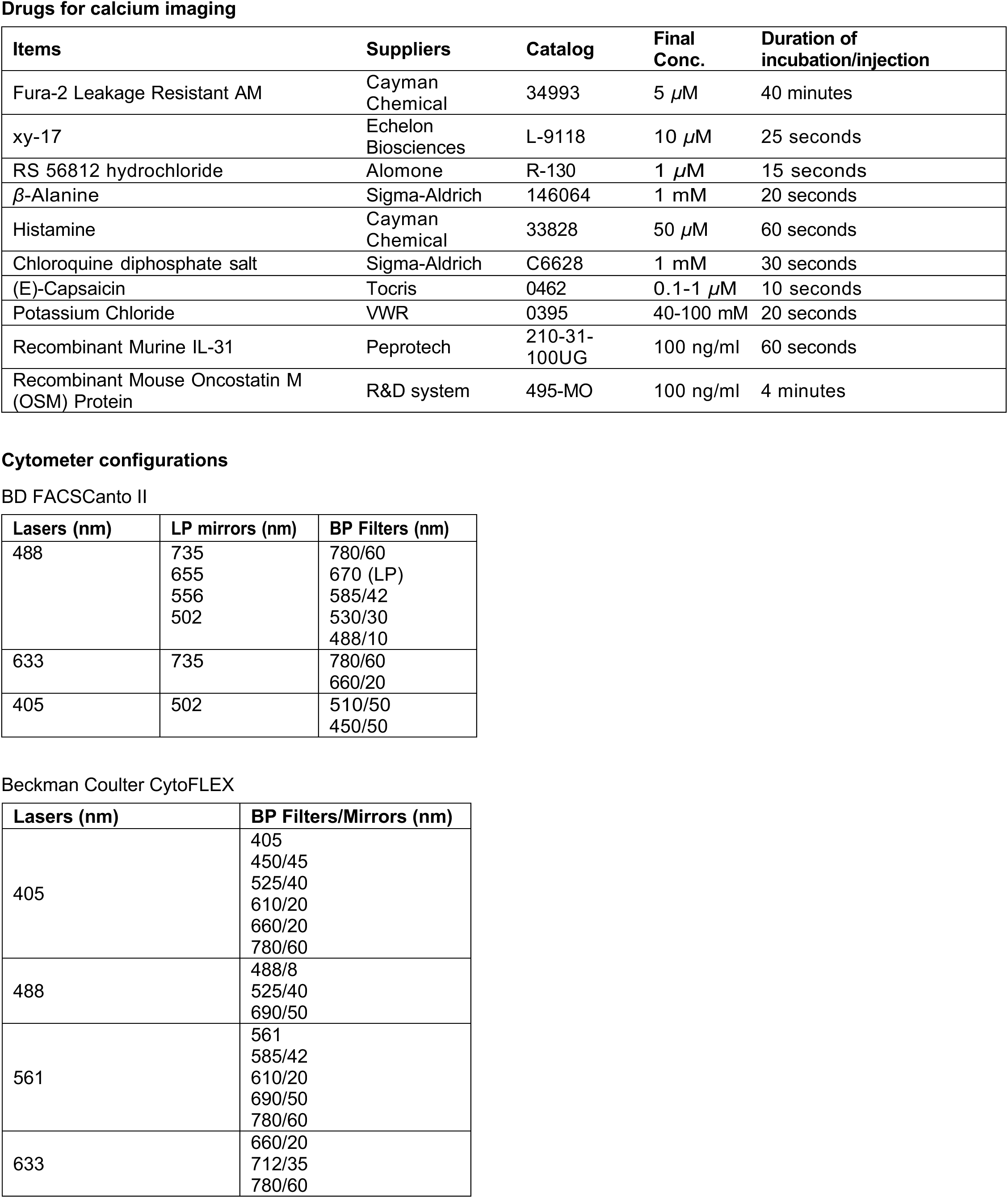
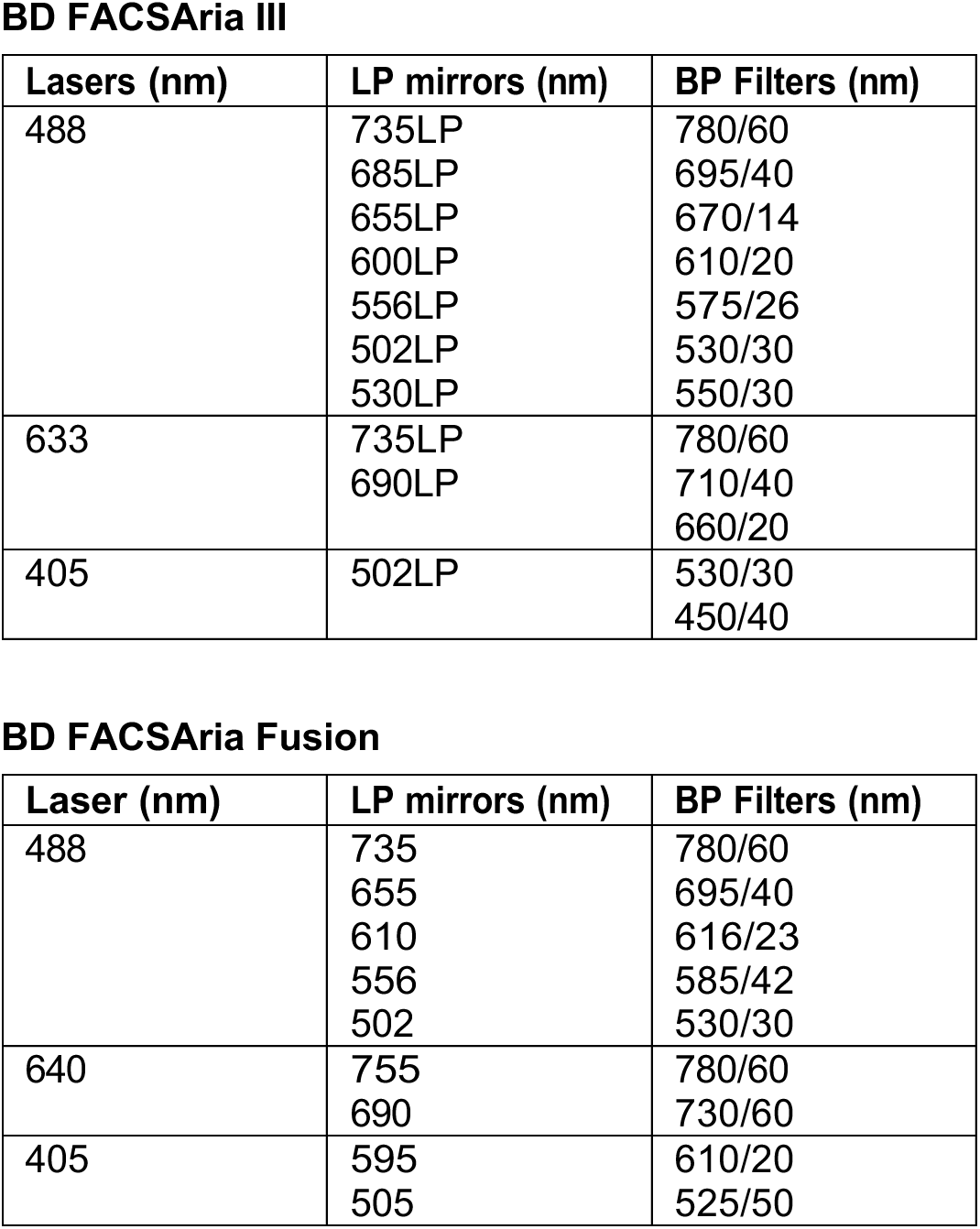

